# Seasonally increasing parasite load is associated with microbiome dysbiosis in wild bumblebees

**DOI:** 10.1101/2023.11.30.569473

**Authors:** Mark G. Young, Josefine Just, Ye Jin Lee, Thomas McMahon, James Gonzalez, Suegene Noh, David R. Angelini

**Author notes:** **Author contributions** - MY, JJ and DRA designed the experiments. - MY conducted the main analysis, creation of draft figures, and writing of the main text. - SN contributed to data analysis and manuscript editing - DRA secured funding, produced some figure components and was involved in manuscript editing for publication. - MY, JJ, YJL, TM, JG collected specimens and were involved in preliminary data analyses.

## Abstract

The microbiome is increasingly recognized for its complex relationship with host fitness. Bumblebees are host to a characteristic gut microbiome community that is derived and reinforced through social contact between individuals. The bumblebee microbiome is species-poor, and primarily composed from a small number of core taxa that are associated with the greater tribe of corbiculate bees. Experimental findings support a role for the core bumblebee microbiome in resistance to severe infections by a common trypanosomal parasite, *Crithidia bombi*. However, most studies have been small in scale, often considering just one or two bumblebee species, or making use of commercially-reared bees. To better understand the microbiome diversity of wild populations, we have deeply sampled field populations of ten sympatric species found throughout central and down east Maine in a three-year microbiome field survey. We have used 16S amplicon sequencing to produce microbiome community profiles, and qPCR to screen samples for infections by *Crithidia bombi*. The breadth of our dataset has enabled us to test for seasonal and interspecific trends in the microbiome community. Controlling for these external sources of variation, we have identified microbial factors associated with infection and parasite load that support the role of the core microbiome in resistance to severe infection.

## Introduction

Bumblebees are host to a characteristic gut microbiome community. As eusocial corbiculate bees (alongside honey bees and stingless bees), bumblebees are primarily colonized by the small set of corbiculate core genera, *Snodgrassella, Gilliamella, Bifidobacterium,* and *Lactobacillus* (Firm-4 and Firm-5) ^1–3^. Evolution of eusocial behavior was likely intertwined with the establishment of the corbiculate core microbiome, as the core taxa do not colonize related non-eusocial bees ^3^. Within colonies, social behavior is the main driver of the homogenization of microbiome communities. Newly eclosed bees must be raised alongside other adults, or their feces, to establish the characteristic microbiome ^4,5^. This is in contrast to some other insects, which possess specialized bacteriocyte cells and receive their bacterial endosymbionts through direct vertical inheritance. Between colonies, foraging provides a route for the transmission of microbes ^6^. Pollinators deposit microbes on flowers ^7^, and gut-adapted bacteria are able to persist on flowers ^8^. Conspecific microbial transmission is likely common as bumblebees are able to discriminate between flowers, showing preference for specific floral rewards ^9–11^.

The bumblebee microbiome is distinct from that of other eusocial bees despite the potential for microbiota transmission between sympatric bee genera. Besides the corbiculate core microbiota, bumblebees have symbioses with a unique set of taxa, including *Lactobacillus* Firm-3, *Bombiscardovia*, and *Candidatus Schmidhempelia*^12^. Compositionally, bumblebee, honey bee, and stingless bee microbiomes are unique and dissimilar in proportion to the phylogenetic divergence between their hosts ^12^. Reflecting host specificity, the strain-level phylogenies of two bee endosymbionts, *Gilliamella* and *Snodgrassella*, correlate more strongly with the phylogenies of their hosts than with their hosts’ geographic ranges ^12,13^. Although strains of *Snodgrassella* specific to *Apis* and *Bombus* hosts are able to colonize non-native hosts under laboratory conditions, they are unsuccessful at achieving high abundances and are rapidly out-competed by native strains, even at numerical advantages of 10:1^14^.

Parasitism is a focal point for bumblebee conservation and agricultural bee husbandry. Throughout the 20th century, bumblebee species throughout Europe and North America have contracted in range, a sign of continuing decline in colony survival ^15–19^. Parasites do not pose a novel threat to bumblebees, and wild bees successfully mount immune responses against a diverse array of protozoa, nematodes, fungi, mites, prokaryotes, and viruses ^20,21^. However, synergy with other stressors exacerbates parasite virulence, causing outsized impacts on colony survival. This “context-dependent virulence” ^22^ is exemplified by a trypanosome endemic to Europe and North America, *Crithidia bombi*. Where *Crithidia* is endemic colony infection rates can reach 80 to 100% by the end of summer ^23–26^. The parasite spreads within nests through coprophagy,^27,28^ and between colonies through foraging on the same flowers ^29^. Infections only become severe when hosts are exposed to concurrent stressors. A major source of stress for wild queen bees is hibernation, and infection by *C. bombi* reduces the likelihood of successful hibernation by 40%^22^. Context-dependent virulence can be replicated experimentally. When raised under favorable conditions, *Crithdia*-infected bees show no excess mortality in comparison to non-infected controls. When starved, excess mortality increases to 50% ^30^. Anthropogenic stressors, such as pesticide exposure or habitat loss, likely also drive context-dependent virulence, contributing to modern declines of bumblebee populations.

Increasingly, it is accepted that the gut microbiome influences colony survival through its role in immunity. Organisms in the core bumblebee microbiome form a biofilm along the epithelium of the hindgut^31^ and have genomes enriched for gut colonization factors ^14,5,32,33^. Together, they may form a physical barrier against infections, as well as influence host immune responses. In the context of *Crithidia*, experimental results support a mechanistic role in resistance to severe infection. Workers raised under sterile conditions suffer far greater parasite loads than workers with normal microbial communities ^4^. Microbiome transplants between *Crithidia-*resistant and susceptible hosts have been shown to alter transcription of immune genes, with susceptible hosts adopting resistant-like gene expression after transplant ^34^. In *B. impatiens*, high relative abundances of the core taxa *Apibacter, Lactobacillus* Firm-5, and *Gilliamella* have been shown to inhibit severe infection^35^.

Observational studies indicate that these results may translate to wild populations ^35,36^. Diversity of non-core taxa has been associated with increased infection load ^36^, and the relative abundances of core taxa have been shown to have negative correlations with infection ^4,36^. However, the studies that we have considered have been limited in scale, and poorly equipped to answer questions of inter-species and inter-site heterogeneity of wild populations. They have also disagreed on the significance of relationships between specific core taxa and infection severity, one even finding no association between microbiome and infection at all ^37^. To address this issue, we have conducted a three-year field survey, deeply sampling populations across coastal Maine, and exploring how microbiota change with the landscape. The scale and breadth of our dataset has enabled us to examine trends of microbiome composition with *Crithidia* infection, as well as isolate inter-season, inter-site, and inter-species variation. We found consistent seasonal changes in bumblebee microbiome composition, and identified variation in the relative abundances of core taxa among host species. Controlling for external sources of variation, parasite load had an inverse relationship with the relative abundances of a few core taxa, and infection was characterized by an increase of what appeared to be bacterial opportunists, derived from local environments.

## Results

### Wild bumblebee species share a core microbiome community

We used 16S ribosomal RNA amplicon sequencing to profile the gut microbiome communities of 638 bumblebees collected during the course of a three year field survey of Maine’s wild pollinators. Our sample set included workers (n=505), queens (n=45), and males (n=88) of ten sympatric bumblebee species from 64 ecologically diverse sites (**Figure 1A**). We did not visit all sites evenly; some were longitudinally sampled during each year of the field study, while others were visited opportunistically, or just once (**Figure 1B**). We targeted the v6-v8 region of 16S. Samples had a mean depth of 51,927 paired-end reads after quality control.

**Figure 1:**
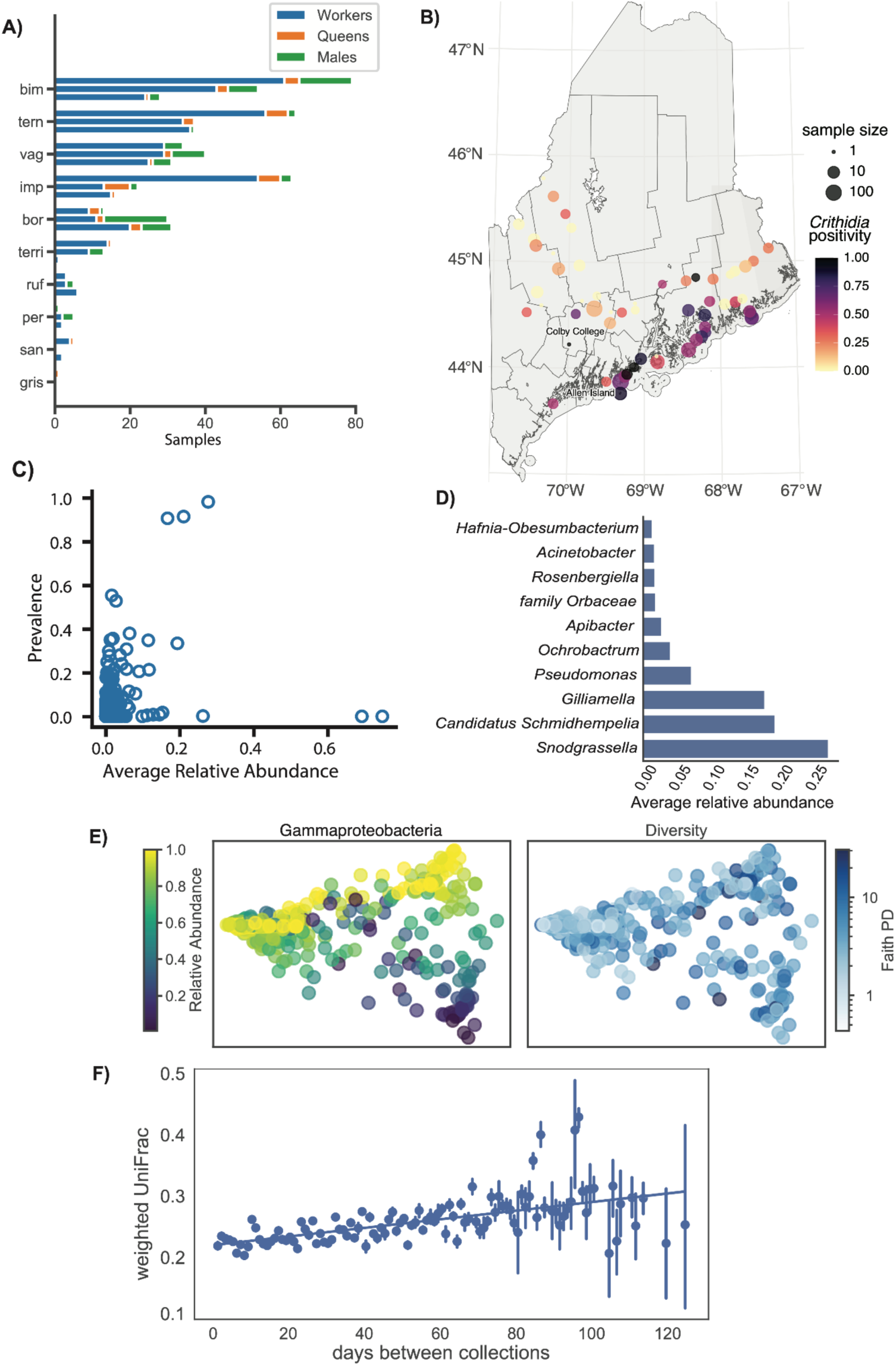
Wild bumblebees share a genus specific microbiome community. A) Wild species and castes are unequally represented in the field survey dataset. The prevalence of species is represented by horizontal bars, broken down by caste (color) and year (vertical stacks). B) Map of Maine with markers for collection sites, sized by number of samples collected per site, and colored by *Crithidia* infection prevalence. C) Prevalence across samples versus average relative abundance distribution of OTUs. The plotted value for average relative abundance is the average non-zero relative abundance. D) Average relative abundance of most frequently identified microbial genera across the full dataset, including samples where they were not detected. E) PCoA plot of weighted UniFrac diversity, shaded by Gammaproteobacteria relative abundance (left) and Faith’s phylogenetic diversity (right). Gammaproteobacteria relative abundance and Faith’s PD are inversely correlated (Spearman’s *r* = -0.39). F) The difference in microbiome composition between samples was dependent on the difference in their dates of collection (Spearman’s r = 0.10, Mantel test, p < 0.001). Difference in collection date was calculated from day-of-year dates, in order to test for trends consistent across the years of the field survey.

Consistent with prior studies, a small number of taxa accounted for nearly all of the bumblebee microbiome. Of 2388 OTU99s detected within our sample set, just 82 were observed in multiple samples at an average relative abundance greater than 1% (**Figure 1C**; **Supplementary Table 1**). Samples were overwhelmingly colonized by OTUs classified as Gammaproteobacteria (median 94% relative abundance), and a small number of genera previously annotated as being part of the bumblebee core microbiome (**Table 1**), three of which (*Snodgrassella, Candidatus Schmidhempelia,* and *Gilliamella*) accounted for 65% average composition (**Figure 1D**). Samples with lower relative abundance of Gammaproteobacteria were characterized by higher Faith’s phylogenetic diversity (**Figure 1E**; spearman *r* = -0.39).

**Table 1:**
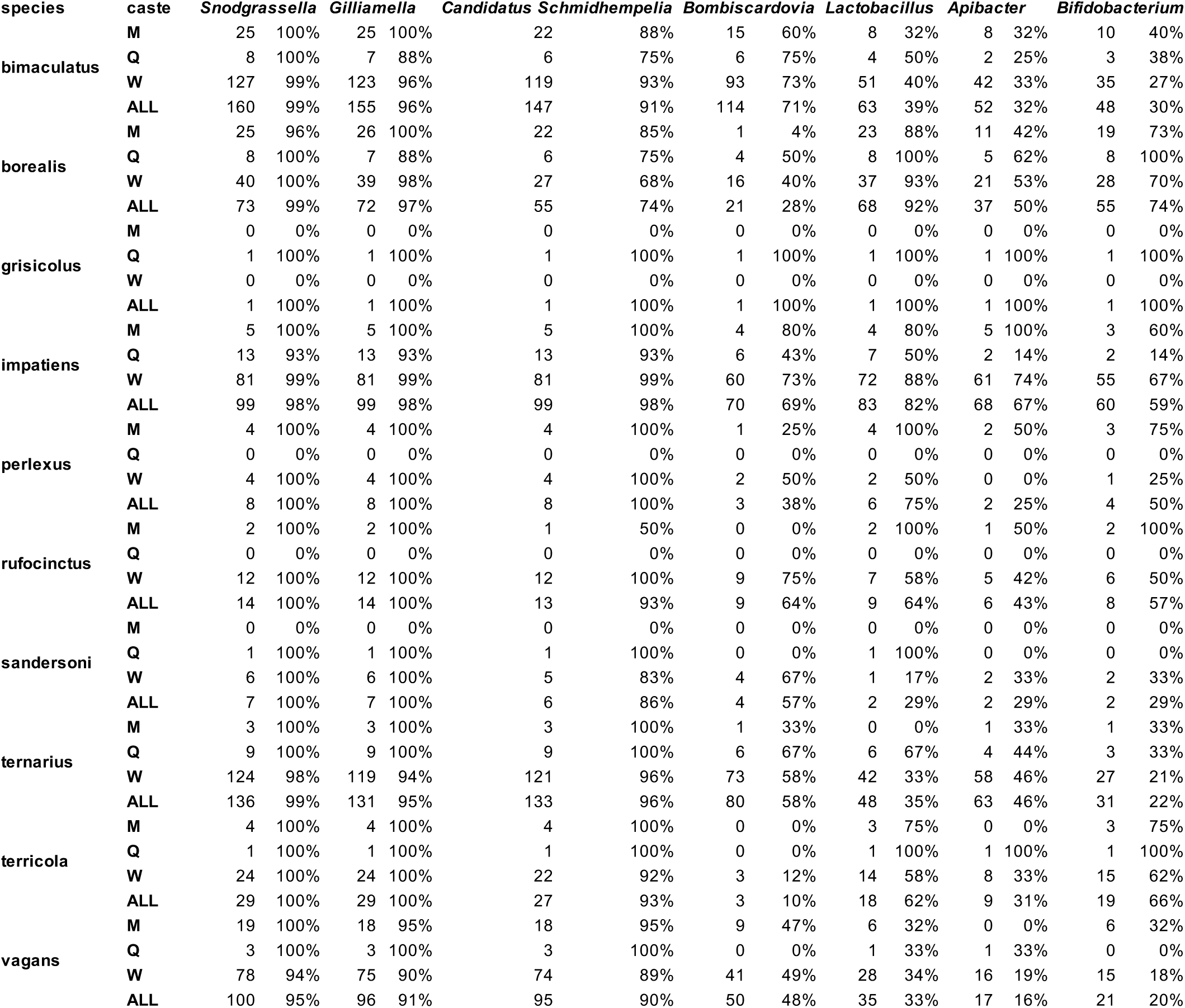
Prevalence of core microbes by host species.

We tested for stratification of microbiome structure by host species, host caste, collection site, and collection month with marginal permanova. All factors except for host caste explained significant variation in the weighted UniFrac dissimilarity between samples, after multiple hypothesis correction (p<0.05; **Table 2**). However, the size of the effects were small, and samples did not show obvious clustering on principal coordinate plots (**Supplementary Figure 1**). To test for monotonicity, we examined correlations between *β*-diversity and the phylogenetic dissimilarity (sum of branch lengths) between host species, the distance between sites, and differences in collection dates (day-of-year) with Mantel tests (**Table 3**). The relationship between *β*-diversity and days between collection was significant (spearman r=0.10, p < 0.001, **Figure 1F**), indicating bumblebee gut microbiota changed consistently across each summer.

**Table 2:** Weighted UniFrac marginal permanova (adonis2)

| **factor** | **df** | **sum of squares** | **R^2** | **F-score** | **p-value** | **Bonferroni corrected** |
| --- | --- | --- | --- | --- | --- | --- |
| species | 10 | 1.863 | 0.07913 | 6.6381 | 0.001 | 0.004 |
| caste | 2 | 0.1078 | 0.00458 | 1.921 | 0.043 | 0.172 |
| site | 89 | 4.2785 | 0.18172 | 1.7129 | 0.001 | 0.004 |
| month | 4 | 0.4305 | 0.01829 | 3.8352 | 0.001 | 0.004 |
| residual | 532 | 14.9306 | 0.63416 |  |  |  |
| total | 637 | 637 | 23.5439 |  |  |  |

**Table 3:** Weighted UniFrac Mantel Correlation.

| **factor** | **correlation** | **p-value** | **samples** |
| --- | --- | --- | --- |
| **phylogeny** | 0.00136 | 0.961 | 638 |
| **geography** | -0.00611 | 0.753 | 601 |
| **day** | 0.10257 | 0.001 | 638 |

### The phylogenetic divergence of *Bifidobacterium, Saccharibacter,* and *Gilliamella* correlate with those of their hosts

Many of the microbiome taxa identified in the field survey dataset were composed of multiple unique OTUs, including each of the 28 taxa that were fully classified to the genus level and were at least moderately prevalent (>10% of samples) (**Supplementary** Figure 2). Potentially OTUs of the same genus corresponded to species or strain-level diversity. We quantified the relative abundance-weighted phylogenetic dissimilarity between the genus-specific fractions of samples with weighted UniFrac. For the core genera *Bifidobacterium* and *Gilliamella,* there was a Spearman correlation between dissimilarity in OTU composition between samples and host phylogenetic divergence, even after controlling for inter-site variation (Mantel test stratified by collection site, p < 0.05, **Supplementary Table 2**). The association was also significant for OTUs classified as belonging to the genus *Saccharibacter*. It is possible that some or all of the OTUs classified as *Saccharibacter* belonged to the closely related core genus *Parasaccharibacter*, which was not included in our classifier. These correlations indicate phylogenetic diversification of endosymbionts alongside their bumblebee hosts, and that for these three genera, host specificity had a greater effect on microbiome composition than local transmission.

### Crithidia infections are common throughout Maine and increase in prevalence over the course of the summer

Hosts from all three summers of the field survey were infected with the parasite *Crithidia bombi*. The distribution of estimated infection loads was bimodal, with a left peak below our predetermined detection threshold (1000 copies/ng), potentially corresponding to low-level, latent infections **(Supplementary Figure 3)**. We consider samples passing our originally defined threshold as positive infections in our analysis.

The overall *C. bombi* infection positivity rate was 45%, but varied significantly between local populations. To compare between locations, we used five sites that had been deeply sampled (≥15 samples/summer) over multiple summers of the field survey: two geographically isolated offshore islands (Vinalhaven and Allen Island), two islands reachable by bridge (Great Wass and Swans Island), and one mainland site (Colby College). The single summer positivity rates were significantly different between sites (ꭓ^2^ test, p < 2.7×10^-12^) and greatest for the island sites (**Figure 1B**). Over 50% of samples collected from Allen Island, Great Wass Island, and Swans Island tested positive, while only 15.6% from Colby College were positive for *Crithidia*.

Infection positivity rate also consistently increased over the course of each summer. We used a generalized linear mixed effects model to measure the dependence of infection status on host collection date and caste, with random effects to capture inter-site and inter-species variation **(Supplementary Table 3)**. There was no difference in infection positivity rate between castes, but the odds of infection increased by 2.6% (p < 0.001) for each day of the summer.Visually, the increase appeared to continue until collections ended in early September (**Figure 2A**), at odds with midsummer peaks in infection rate reported in Popp et al. (2012) ^38^. Consistent with the inter-site comparison of infection rates above, the site random effect explained a large and significant proportion of total variation (likelihood ratio test; p < 1×10^-12^). To capture inter-species differences, we used independent and identically distributed random intercepts (i.i.d.), as well as a correlation structure proportional to the phylogenetic dissimilarity between hosts. Only the i.i.d. encoding explained significant variation (LRT; p < 0.0008), indicating that the differences in infection rate between species were driven by something other than their phylogenetic relationships. The variation explained by species was relatively small, roughly 14% of that explained by differences in site (**Supplementary Table 3**).

**Figure 2:**
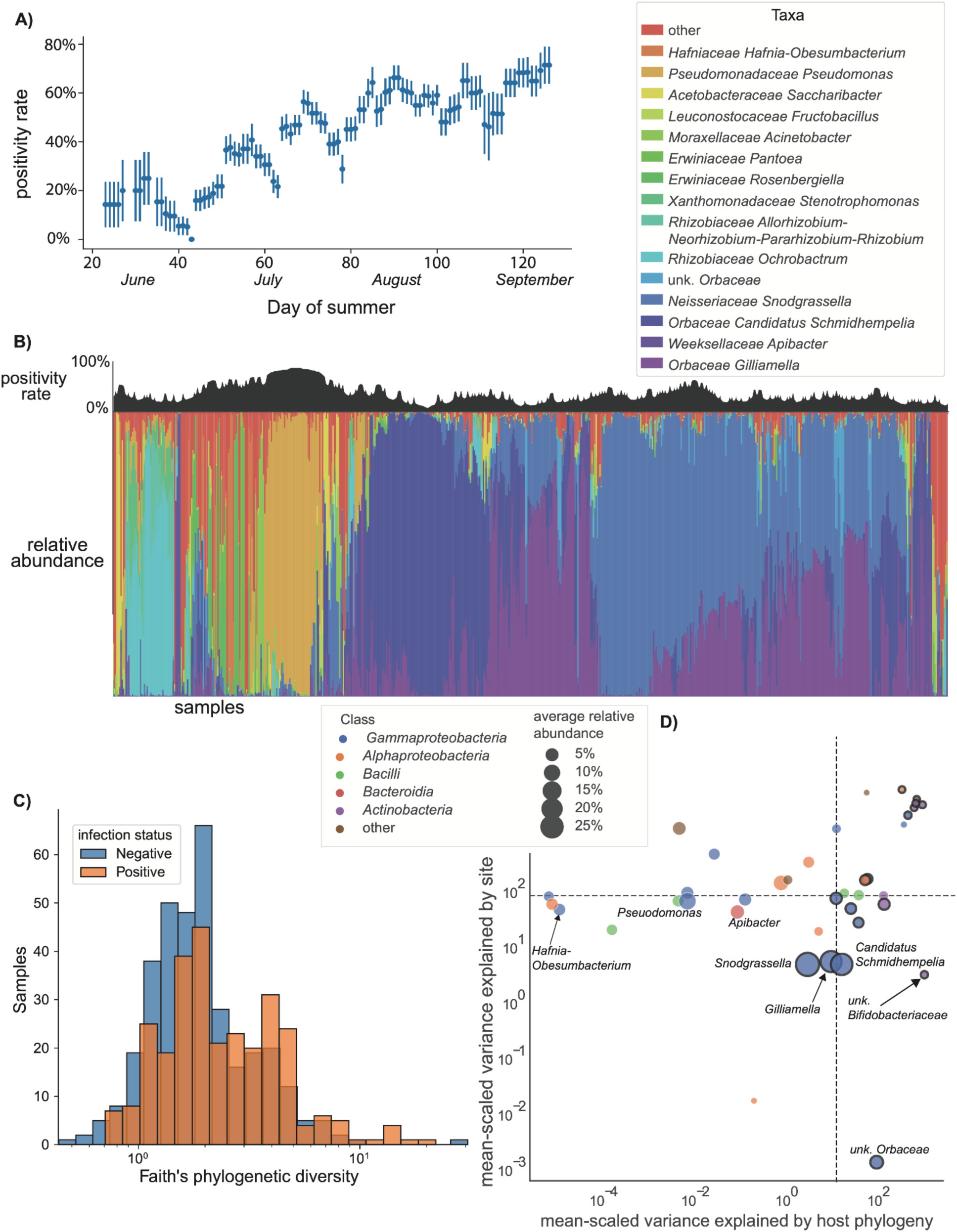
Bumblebee microbiome community is associated with host ecology and *Crithidia* infection status. A) *Crithdia* infection rate increases over the course of the summer. Positivity rates were estimated as the proportion of positive samples collected within five days before or after a given day. Rate estimates include samples from all three years of the field survey. Standard deviations of estimates are shown as vertical bars. B) *Crithidia* infections are associated with microbiome dysbiosis. The relative abundances of the 15 most abundant microbial taxa are shown for all 638 samples. Samples are sorted along the horizontal axis by agglomerative clustering on weighted unifrac distance. *Crithidia* positivity rate was estimated using a sliding window with a radius of 20 samples, and is shown above the taxonomy plot. C) *Crithidia* positive samples had greater Faith’s phylogenetic diversity. Faith’s phylogenetic diversity, a phylogeny-aware alpha diversity metric, was calculated for microbiome communities with qiime2. Horizontal axis is log scaled. D) Linear mixed effects models were used to assess the association between the relative abundances of individual taxa with *Crithidia* infection, collection date, collection site, and host species. Points represent the variance in relative abundance explained by the phylogenetic and site random effects. Mean-scaled variance is plotted. Medians are indicated by dashed lines. Taxa for which significant variation is explained by host phylogeny (ɑ = 0.05) are given black outlines.

### *Crithidia* infections were associated with changes to the relative abundances of a small number of taxa

The relative abundances of a small number of microbial taxa were different between *Crithidia* infected and non-infected (healthy) bees, suggestive of infection-associated dysbiosis. To test for differential abundance, we used linear mixed effects models, as described in Maaslin ^39^, with fixed and random effects to control for the effects of host species, collection date, and collection site. We tested whether *Crithidia* infection was associated with the 38 genus-level and 9 class-level taxonomic bins found across 10% or more samples, and an ‘other’ bin of less prevalent or unassigned taxa.The ‘other’ bins contained highly diverse assemblages of low abundance taxa. The genus level bin contained 620 unique taxa, representative of 2.2% of average relative abundance. The class level bin contained 74 groupings, for 0.7% of average relative abundance. The genera *Pseudomonas* and *Hafnia-Obesumbacterium,* as well as the class-level ‘other’ bin had significant positive associations with infection (**Supplementary Table 4**). Though at a weaker significance (p < 0.10), the class Gammaproteobacteria was positively associated with infection, while the genera *Apibacter* and *Gilliamella* and class Bacteroidia were negatively associated with infection. Qualitatively, the health-associated taxa appeared to be consistent and prevalent members of the bumblebee microbiome, while besides Gammaproteobacteria, infection-associated taxa were only found at high relative abundance in small clusters of primarily *Crithidia*-positive samples (**Figure 2B**). Infected samples also had greater Faith’s phylogenetic diversity (**Figure 2C**), indicating the presence of additional non-core taxa.

Many taxa changed in relative abundance over the course of the summer, consistent with the compositional changes noted above, but associations with collection date did not discriminate between infection- and health-associated microbes. At the genus level, the infection-associated *Hafnia-Obesumbacterium* and grouping of ‘other’ taxa both increased in relative abundance with collection date, mirroring the date dependent rise in *Crithidia* infections (p< 0.05). However, relative abundances of the health-associated *Apibacter* and *Gilliamella* did as well. For the full microbiome, the effect of collection date was relatively large, but sporadic. Twenty-five of 38 genus level taxa, representative of 42% of the average relative abundance, were significantly associated with date (positive or negative), while only three relatively minor classes were, Bacteroidia, Bacilli, and Cyanobacteria (**Supplementary Table 4**, **Supplementary Figure 4**).

The random effect structures from the differential abundance tests helped to determine the contributions of host specificity and local transmission to the relative abundances of specific taxa. The two taxa with the largest ratio of variance explained by host vs. site were unknown genera of the families Bifidobacterium and Orbaceae (**Figure 2D**), both of which are well adapted to the tribe of corbiculate bees ^3^. Relative abundances of other core taxa, including *Gilliamella*, *Snodgrassella*, and *Candidatus Schmidhempelia*, were also significantly dependent on host phylogeny (likelihood ratio test; p< 0.05), reflecting adaptation to the bumblebee gut. In contrast, neither *Pseudomonas* nor *Hafnia-Obesumbacterium* showed any association with host phylogeny (p = 1), while instead being largely dependent on collection site (p < 1×10^-6^, **Supplementary Table 4,** **Figure 2D**), suggesting they were environmentally derived opportunists, rather than parasites specifically adapted to bumblebee hosts.

### Severe infection is associated with increasingly dysbiotic microbiome composition

Among *Crithidia*-positive samples, there was great heterogeneity in infection severity. Positive samples exhibited a 230-fold range in infection load (**Supplementary Figure 2**). We used linear mixed effects models to assess the relationship between infection severity and the relative abundances of bacterial taxa. We fit models of the same form as described above, with the binary indicator for infection exchanged for log-transformed infection load. We used only the subset of samples passing the positivity threshold.

Across the full set of taxa, there was a strong correlation between the fitted values from the two modeling approaches (Pearson’s r = 0.80), indicating that changes in relative abundances with increased severity largely mirrored the differences between non-infected and infected bees. Severity was associated with an increase in the relative abundance of *Pseudomonas* (p=1x10^-8^), and decreases in *Gilliamella* (p=0.0001) and *Apibacter* (p=0.0002). There were also additional core microbiota that decreased in relative abundance with severity, that were not associated with binary infection status: *Snodgrassella* (p = 0.06), *Candidatus Schmidhempelia* (p = 0.02), and an unknown genus of family *Orbaceae* (p = 0.02) (**Figure 3**; **Supplementary Table 5)**. At the class level, the relative abundance of Alphaproteobacteria increased with infection severity, despite not showing a significant relationship with binary positivity, and the relative abundance of Gammaproteobacteria decreased, despite being slightly greater in infected bees on average. The fold changes associated with Gammaproteobacteria were relatively small, likely reflective of its high average relative abundance in the bumblebee gut. For both differential abundance tests, the bin of ‘other’ taxa had the largest magnitude of association with *Crithidia* infection, reflective of an increase in randomness of community composition accompanying the depletion of core taxa.

**Figure 3:**
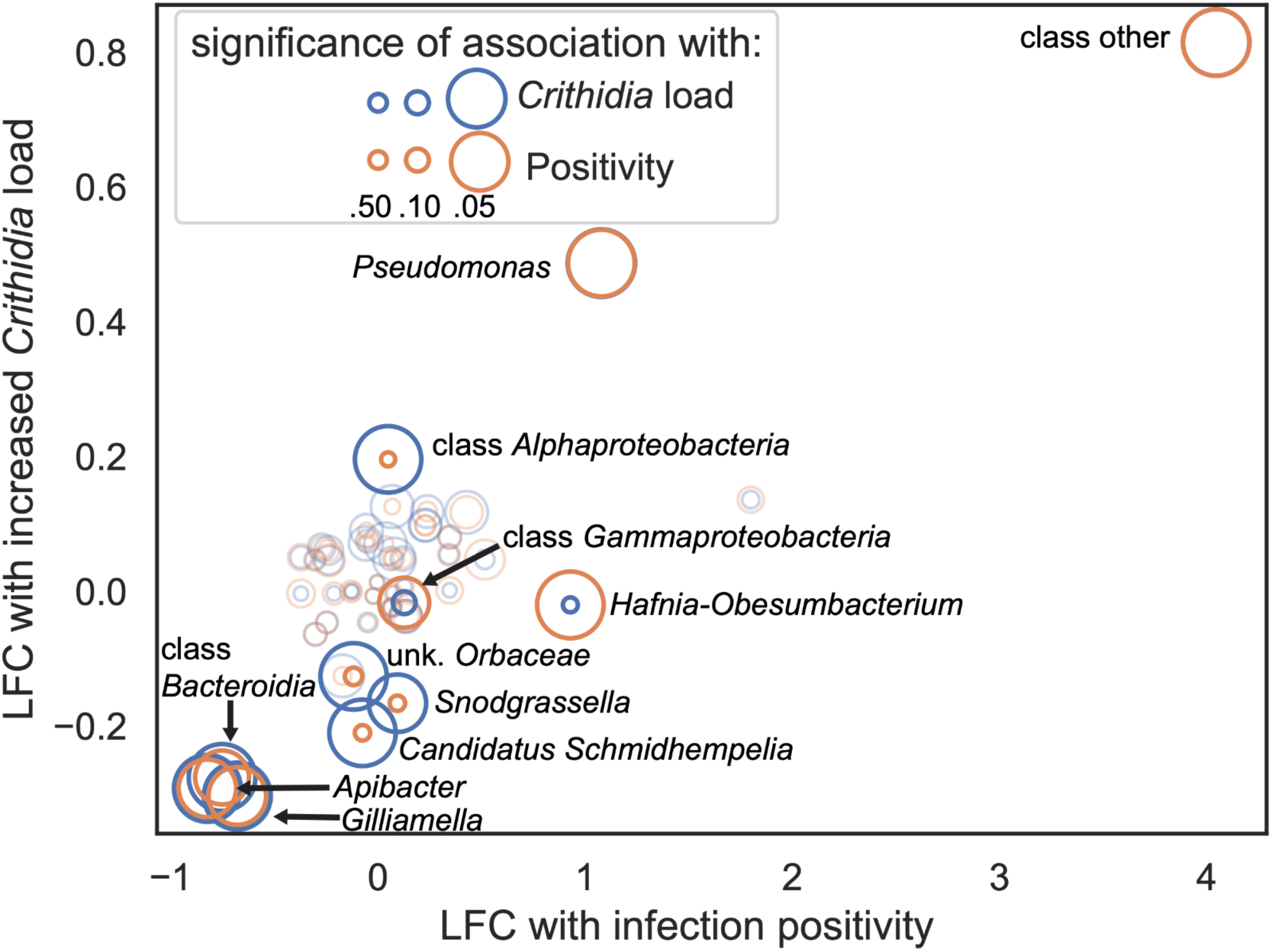
Bacterial taxa associated with infection were also associated with increased infection load. Differential abundance models were used to measure the difference in relative abundance of bacterial taxa between infected and non-infected samples (horizontal axis), as well as to test for changes in relative abundance with increased infection load (vertical axis). Concentric circles represent the FDR-corrected significance of the relationships between taxa relative abundance and infection (orange) and infection load (blue). Larger circles correspond to increased significance. Taxa with weak associations (p<0.1) with either infection load or positivity (or both) are indicated with darker colors. For *Pseudomonas* and ‘other’ the associations had equal significance, causing the circles to overlap. Differential abundance models were fit for both genus and class-level taxa, fitted values for both are included on the plot.

## Discussion

### Evidence for a core bumblebee microbiome

The findings of our Maine bumblebee field survey were consistent with a model of a robust and coevolved bumblebee gut microbiome. Samples from all ten *Bombus* species were primarily colonized by bacterial genera previously annotated as being part of the core bumblebee microbiome, mostly of the class Gammaproteobacteria (**Table 1**; **Figure 1D**). The resolution of our study is limited by our use of short read 16S amplicons, and the taxa represented in the Silva 16S database (release 132.99) we used for taxonomic classification. We were unable to differentiate between phylotypes of the genus *Lactobacillus,* nor identify the genus *Parasaccharibacter*, both considered low abundance members of the core microbiome^12^. In spite of these limitations, we still retained a high degree of taxonomic sensitivity. Across the full dataset, we identified 2,388 unique OTU99s, which corresponded to 659 genus-level taxonomic bins. *Snodgrassella* and *Gilliamella* were the most frequently observed microbes, colonizing just over 98% and 96% of samples, respectively (**Table 1**). Similarly high prevalences of these taxa have been observed in other surveys of field caught bees ^35,36,40^.

The field survey dataset additionally provided evidence for microbiome divergence between sympatric bumblebee species. Prior characterizations of the bumblebee microbiome have focused on the genus as a whole, often contrasted against the other tribes of corbiculate bees. We found that members of the ‘bumblebee core microbiome’ (including *Snodgrassella, Gilliamella,* and *Candidatus Schmidhempelia*) varied in relative abundance between sympatric bumblebee species. The degree of microbiome divergence was proportional to the phylogenetic dissimilarity between hosts (**Figure 2D**). At a more granular level, we also found evidence of host specificity of OTUs of three genera, *Bifidobacterium, Saccharibacter,* and *Gilliamella* (**Supplementary Table 2**). Similar phylogenetic associations have been reported in large-scale comparisons of corbiculate bees ^12^, and confirmed experimentally, with strains isolated from bumblebees and honeybees ^31^. To our knowledge this is the first observation of host specificity within bumblebees. Notably, we did not observe OTU diversity within *Snodgrassella*, with one major OTU accounting for >99.9% of all *Snodgrassella* relative abundance (**Supplementary Figure 2**). This could reflect a below average mutation rate at the 16S loci chosen for identification (V6-V8) or a technical artifact of the choice to cluster ASVs into OTU99s.

### Gut flora change with the seasons

Seasonal variation in bumblebee ecology was reflected in the microbiome. The bumblebee life cycle can be divided into two major phases, colony formation in the summer, and hibernation in the winter. Only the gynes (reproductive females) hibernate, before emerging in the spring to form colonies of their own. Over the winter, core taxa decrease in relative abundance, while the microbiome increases in richness and evenness ^13,41^. In the summer phase of the life cycle, the bumblebee microbiome undergoes selective pressure from diet, and is homogenized between colonies through pollination-mediated exchange of microbes. Consistent with local exchange, we found that the collection site of samples was a significant determinant of their microbiome communities (**Table 2**). We also found date-dependent variation in community structure that was consistent across all three years of the field survey, implicating the consistent annual trends in lifecycle and floral rewards as driving factors (**Figure 1F**; **Table 2**). Despite their different ecological roles, we found no significant inter-caste variation in microbiome composition **(Table 2**) when controlling for confounding effects of collection date, suggesting homogeneity between members of the same colonies.

### *Crithidia* infection is associated with dysbiosis of the gut microbiome

*Crithdia* infections were seasonal and highly prevalent within our dataset. Positivity rate climbed linearly from around 10% in May to near 70% in September, consistent with continual parasite transmission within and between local colonies (**Figure 2A**). Infection positivity rates were also highly variable between sites, possibly as a result of variable colonization by the parasite or population bottlenecks caused by overwintering. In agreement with prior characterizations of context-dependent virulence ^22^ parasite load was bimodally distributed, with separate peaks possibly corresponding to mild and virulent infections (**Supplemental Figure 2**).

Parasite loads greater than our threshold for positivity (1000 copies/ ng gut) were associated with the abundance of non-core taxa. We observed significant increases in the relative abundance of ‘other’ taxa (too low-abundance for individual testing), as well as an increase in alpha diversity with infection (**Figure 2**), recapitulating prior findings of associations between the non-core microbiome and *Crithidia* ^35,36,42^. However, we did not find any strong candidates for indicator species for infection. The two taxa with increased relative abundance in infected samples, *Pseudomonas* and *Hafnia-Obesumbacterium*, varied significantly between collection sites, and were not consistent indicators of infection (**Figure 2D**).

Changes to the core microbiome were significant in the contrast between weak and strong infections. Core microbes *Gilliamella, Apibacter, Candidatus Schmidhempelia,* and an unknown genus of the family *Orbaceae* were significantly negatively associated with parasite load. (**Figure 3**). These changes were consistent with experimental support for the role of the core microbiome in resistance to virulent infection ^4,34^. However, the lack of strong associations with positivity suggests that the core microbiome does little to protect against the spread of non-severe infection.

## Conclusion

Here, we report the results of a three year field survey of the gut microbiome dynamics of wild bumblebees, in the context of infection by an endemic trypanasome, *Crithidia bombi*. We found that the relative abundances of core taxa were inversely related with infection severity, and that non-core microbial taxa were at increased relative abundance in infected samples. Additionally, we found evidence for diversification in the microbiome among sympatric bumblebee species, and patterns of seasonal variation in microbiome community structure that was consistent across all three years. These results are evidence of strong associations between bumblebees and their endosymbionts, and the contribution of the microbiome community to resistance to severe parasitism.

## Materials and Methods

### Sample collection

Foraging bumblebees were collected throughout the state of Maine during summer 2017, 2018, and 2019. Individuals were photographed at the time of capture and stored at -80°C before dissection to remove the gut. Species and caste were confirmed with reference to Williams et al. (2014) ^43^. Maxwell 16 DNA purification kits (Promega, Madison, WI) were used for DNA extraction from gut samples. DNA extractions were performed separately after each field season. Extracted DNA was stored at -20°C prior to sequencing.

### Screening and quantification of *Crithidia bombi*

We screened bumble bee gut DNA samples for *Crithidia bombi* infections with quantitative PCR on a CFX96 Touch real-time thermocycler using iTaq Universal SYBR Green Supermix (BioRad, Hercules, CA) and primers for *C. bombi GAPDH* (Cb’gapdh-52F GCGTACCAGATGAAGTTTGATACG; Cb’gapdh-147R AAGCACATCCGGCTTCTTCA). We used 1000 copies/ng DNA as a threshold for positivity. Samples were initially screened in batches, in order to reduce the total number of reactions. We used a batch size of four, and ran batches in duplicate. We then individually ran samples from positive batches, in triplicate.

### Microbiome sequencing

DNA samples were shipped on ice packs to the Centre for Comparative Genomics and Evolutionary Bioinformatics at Dalhousie University (Halifax, Canada). Samples were used to prepare paired-end 2 × 300-bp MiSeq libraries, with PCR amplification targeting variable regions 6-8 of the bacterial 16S ribosomal RNA gene ^44^. Of 732 bumble bees collected, 638 samples passed standard benchmarks for library quality (**Supplementary Table 6**).

### OTU Clustering

We processed the demultiplexed amplicon sequencing data with QIIME2 (v. 2019.4)^45^. We trimmed the first 34 bp from each read and truncated forward and reverse mates to 280 and 240 bp, respectively, before forming ASVs and removing chimeras with DADA2 (v. 1.1.0)^46^. We then clustered ASVs into OTU99s with VSEARCH (v. 2.21.1)^47^, to reduce what was seen as unnecessary dataset complexity. The ASV table produced by DADA2 contained 3,752 total ASVs, including 2,766 that were only found in a single sample (74%). On average, individual ASVs were only found across 4.77 of 638 samples. Clustering to 99% identity with VSEARCH reduced the dimensionality to 2,388 features. After clustering, a similar proportion of features were found in single samples, (1,823 ASVs, 76%), and the average prevalence of individual features was slightly greater (5.69 samples/feature). Because inter-year and inter-site trends in microbiome diversity were the primary focus of our analysis, we were comfortable with the potential loss of some specificity provided by the unclustered ASV sequences. Since we performed the secondary clustering, we refer to features as OTUs throughout the text.

### Taxonomic classification

We classified OTU representative sequences with a Naive Bayes classifier trained on the SILVA 138 SSU NR database ^48^. We used taxonomic labels with minimum posterior probabilities of 0.7 and classified OTUs to the species level when possible. The Silva database does not contain representative sequences for the different *Lactobacillus* phylotypes, nor the genus *Parasaccaribacter*, which are core members of the *Bombus* microbiome.

### Host filtering

Four OTUs were assigned to the order Hymenoptera by the taxonomic classifier. We confirmed the classification with BLASTn alignments to the NCBI NR database. At least one of the four OTUs was present in 453 of 638 samples (71%). The mean relative abundance of the Hymenoptera OTUs was 0.16%, and varied widely between samples (**Supplementary Table 1**). We removed these OTU from the feature table before diversity metric calculation and differential abundance testing.

### Diversity calculation

We used the phylogeny-aware weighted UniFrac distance for *β*-diversity analyses. To build a *de-novo* phylogeny, we created a multiple sequence alignment of all OTUs with MAFFT (v. 7.505)^49^ and masked positions with conservation less than 40%. We then created an unrooted tree with FastTree2 ^50^ and midpoint-rooted this tree. UniFrac distance calculations were implemented in QIIME2. For ɑ-diversity, we calculated Faith’s phylogenetic diversity, also using the rooted tree and a feature table down-sampled to a depth of 4,000 OTUs, which included 631 (98.9%) of 638 samples.

### Host metadata

We measured the dependence of microbiome composition on host species, collection site, and collection date. To quantify the phylogenetic dissimilarity between bumblebee species, we used the sum of branch lengths from the bumblebee phylogeny reported in Cameron and Hines (2007)^51^. To quantify the difference in collection site between samples, we used the distance in kilometers between individual bumblebee collection sites, and also categorically encoded collection sites. Categorical sites had maximum radii of 5 kilometers, which has been reported to be the maximum foraging range of worker bees^20^. We used day-of-year encoding for collection dates, as well as categorical offsets for each year of the field survey.

### Statistical analysis

All statistical analyses were performed in R v. 4.1.3 ^52^. We used linear models to assess variation in *Crithidia* infection rate, and to test for differential abundance of bacterial taxa. For both applications, we fit models with the *pglmm* function from the R package *phyr* ^53^. We used *pglmm* for its support of generalized linear mixed effects modeling with phylogenetic random effects. For modeling variation in *Crithidia* infection rates, we fit the model:

*Crithdia*∼*days*_*Since*_*May*1 + *caste* + (1|*species*_) + (1|*site*_*year*), with a binomial link function. The variables were defined as:

*Crithdia:* Infection status (positive/negative) of individual samples

*days_Since_May1*: The year-independent collection date of samples, measured as the the days between sample collection and May 1st of the same year.

*caste:* sample caste (male/worker/queen)

*(1|species )*: The phylogenetic effect, consisting of independent and identically distributed intercepts for bumblebee species, as well as a covariance structure proportional to the phylogenetic dissimilarity between species (sum of branch lengths).

*(1|site_year)*: Independent and identically distributed random intercepts for collection sites. Collection sites were encoded separately for each year (ex: AllenIsland_2017, AllenIsland_2018, AllenIsland_2019), in order to capture year-specific variation.

Prior to log-transformation of relative abundances, we additively smoothed zero values with a pseudocount equal to half the smallest non-zero relative abundance. We used Benjamini-Hochberg FDR-corrected p-values to assess significance of associations. We corrected for false discovery rate separately for the genus and class-level models. For all modeling applications, we used likelihood ratio tests (implemented in *phyr*) to quantify the significance of the variation explained by random effects. In the discussion of differential abundance testing results, by ‘species random effect’, we refer to the covariance structure proportional to phylogenetic dissimilarities between host species. All figures were created with Python 3.8.16 (*seaborn* v.0.12.2*, matplotlib* v.3.6.0).

## Data availability

Code and data involved in this analysis are available online at https://github.com/aphanotus/bombus.landscape

## Supporting information

All Tables

## Acknowledgements

Research reported in this publication was supported by Colby College, by an Institutional Development Award (IDeA) from the National Institute of General Medical Sciences of the National Institutes of Health under grant number P20GM0103423, and by grant IOS-1350207 from the National Science Foundation to DRA.

---

**Supplementary Figure 1:**
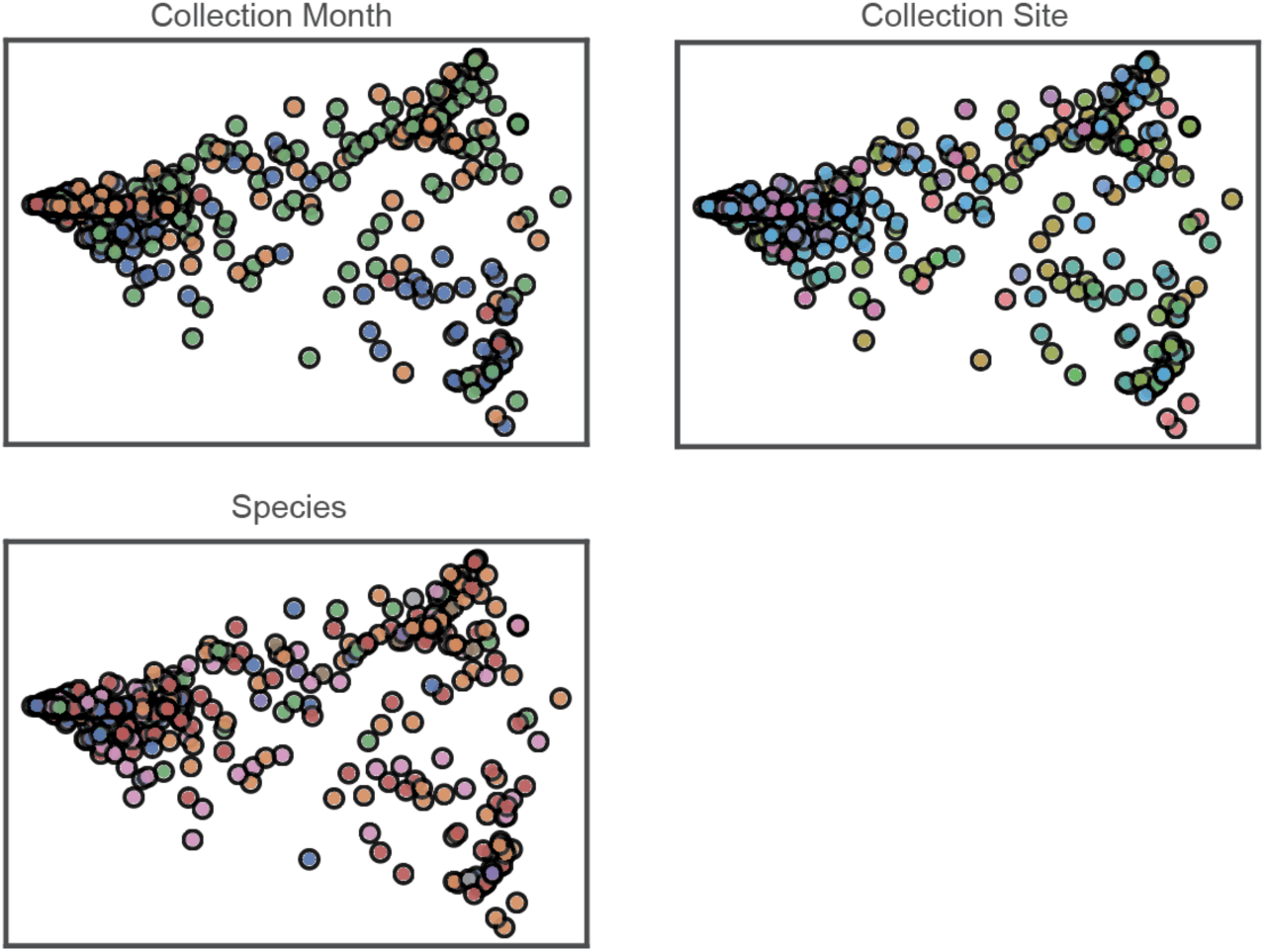
Weighted UniFrac PcoA. Collection month, collection site, and species were all significant determinants of bumblebee microbiome composition (permanova, p < 0.05). However, the size of the effects were small, and samples did not visibly cluster. Points on PcoA plots are colored by categorical metadata (month, site, species). Legends are withheld for length, and because samples do not cluster.

**Supplementary Figure 2:**
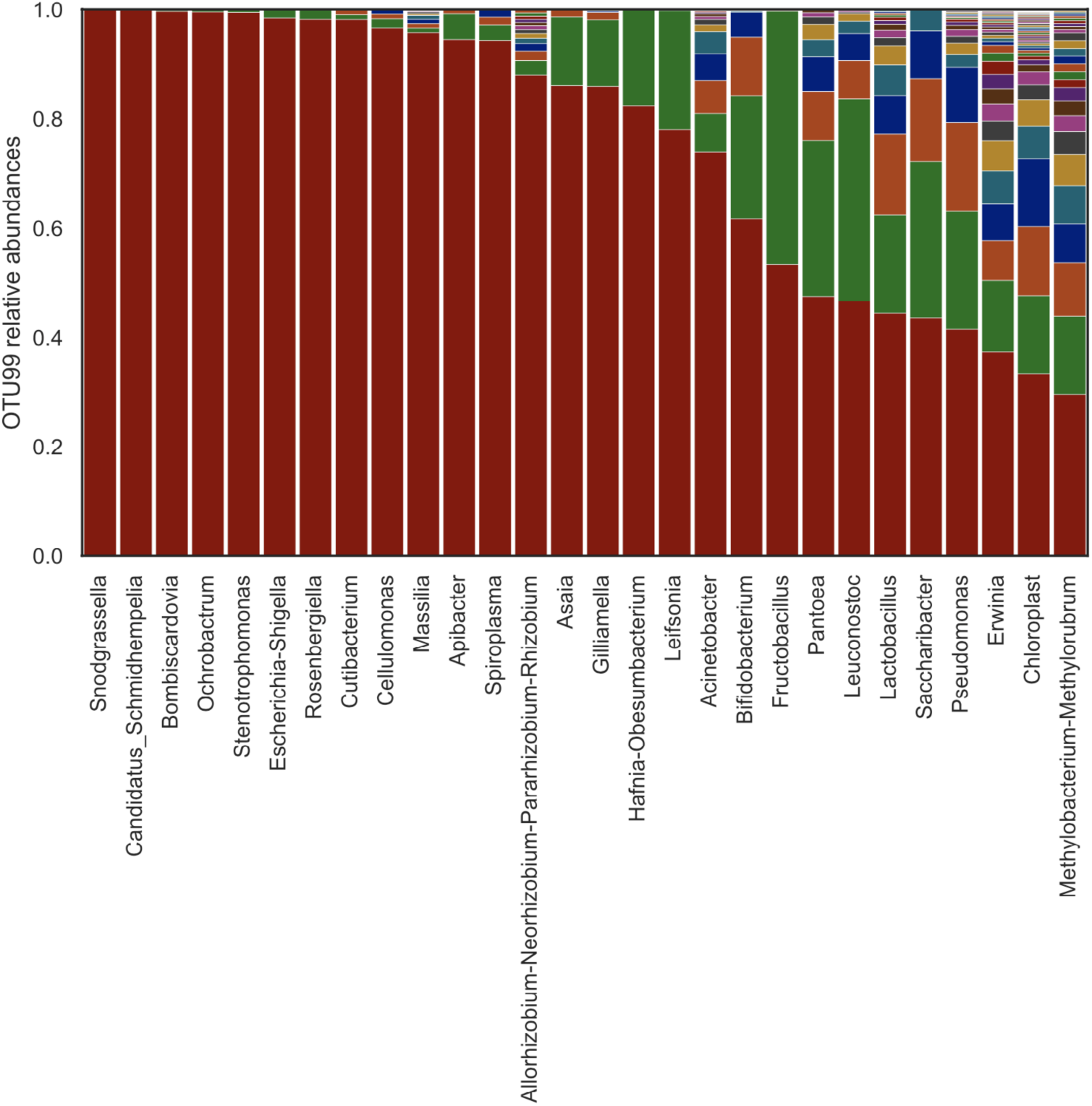
Bacterial genera were composed of multiple OTUs. We selected operational taxonomic units by clustering ASVs from DADA2 into OTU99s with Vsearch. There were 28 clusters of OTUs that were able to be fully classified to the genus level and were at least moderately prevalent in our dataset, being found across 10% or more samples. Colored vertical bars are proportional to the average relative abundances of individual OTUs within the 28 genuses. OTUs are sorted vertically by relative abundance.

**Supplementary Figure 3:**
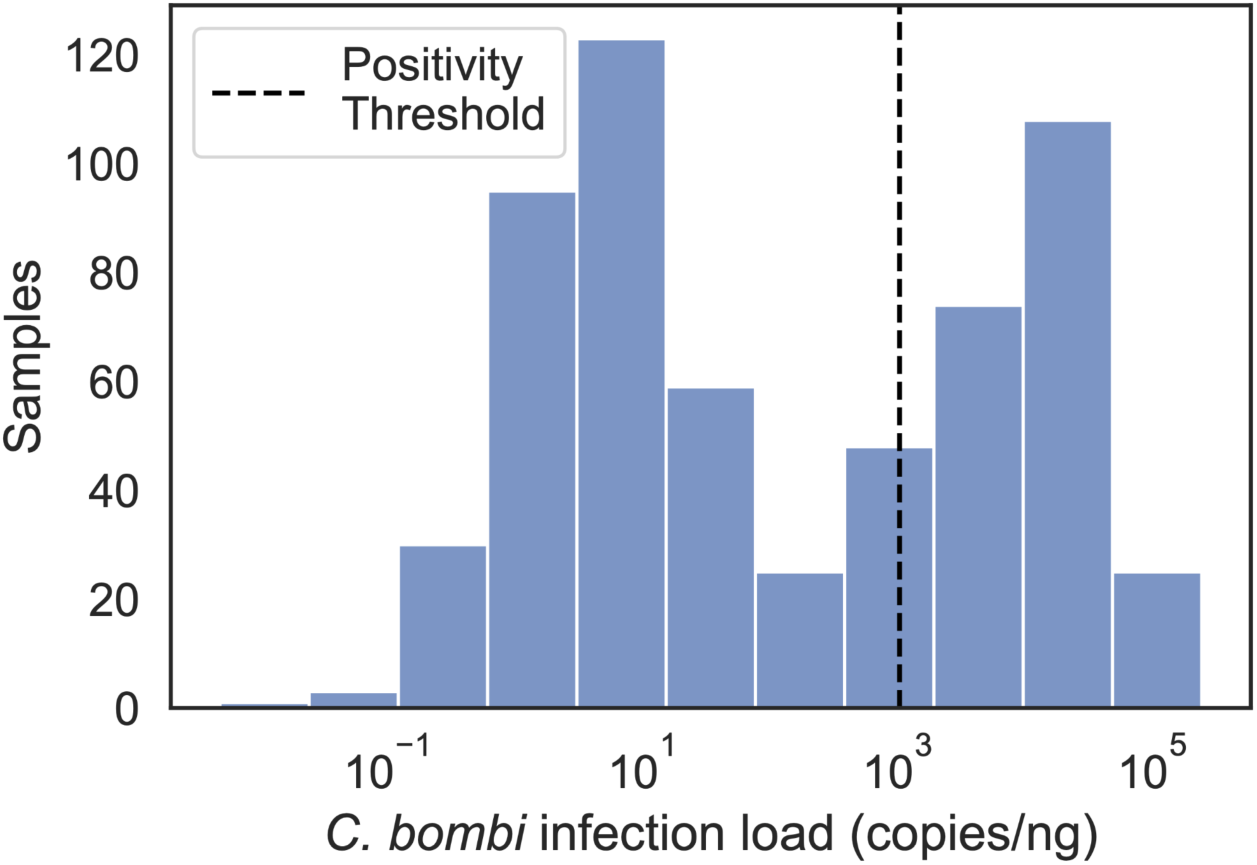
Distribution of *C. bombi* infection load. *Crithidia bombi* infection load was measured with qPCR for *C. bombi GAPDH*. Samples were initially screened with a batched protocol (see Supplementary Methods). Only samples from positive batches were screened individually. We used 1000 copies per ng (dashed vertical bar) as the threshold for positivity. The infection loads of samples run individually were bimodally distributed, with a left peak below the threshold.

**Supplementary Figure 4:**
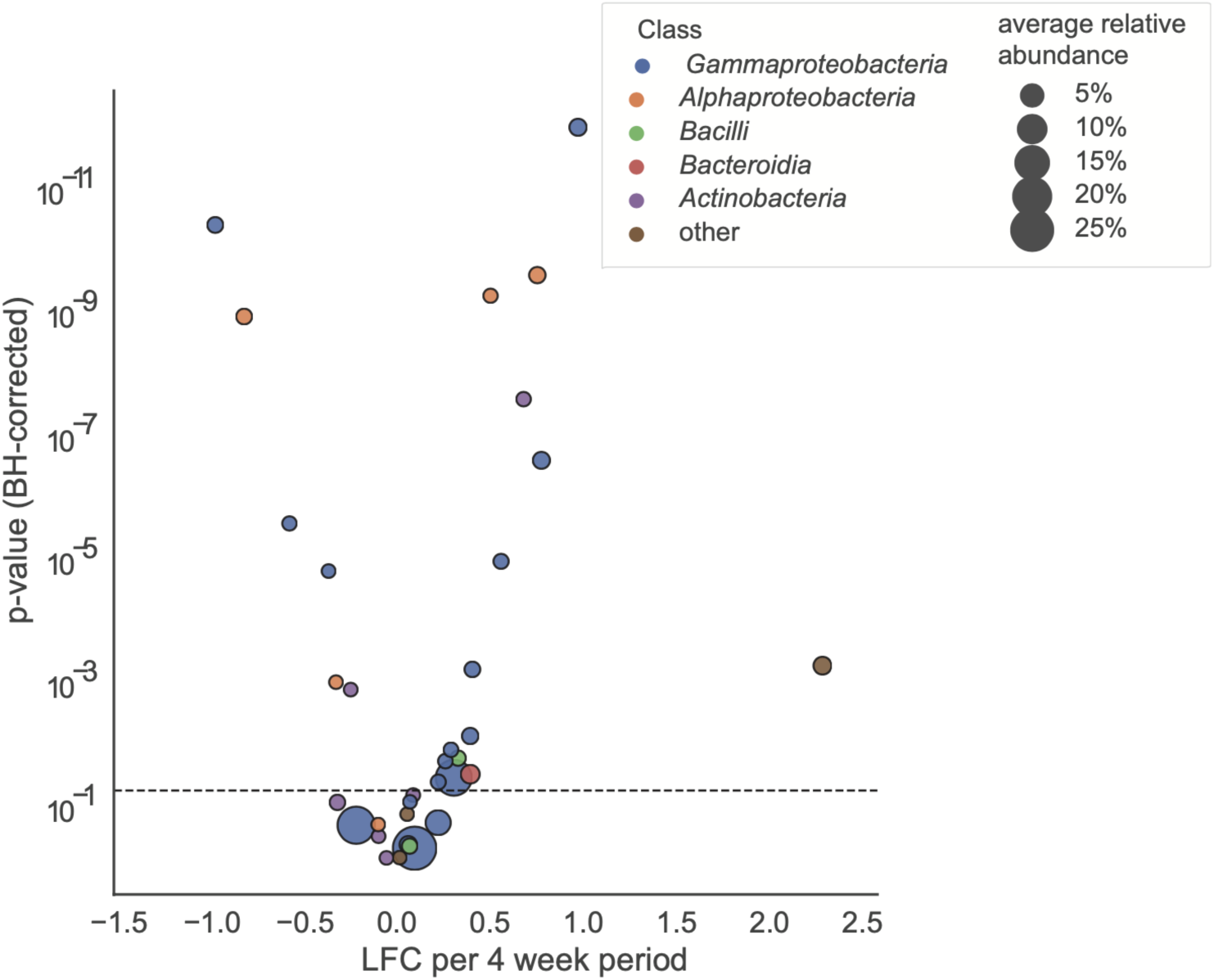
Relative abundances of genera were associated with collection date. Points represent specific genera, colored by class and sized by average relative abundance. The chosen significance threshold (α = 0.05) is indicated by a dashed horizontal bar. The brown ‘other’ point corresponds to a grouping of taxa with insufficient prevalence for individual differential abundance testing.

## Notes

https://github.com/aphanotus/bombus.landscape

### Competing Interest Statement

The authors have declared no competing interest.

## Works Cited

1. Koch, H. & Schmid-Hempel, P. Bacterial Communities in Central European Bumblebees: Low Diversity and High Specificity. Microb. Ecol. 62, 121–133 (2011).

2. Martinson, V. G. et al. A simple and distinctive microbiota associated with honey bees and bumble bees. Mol. Ecol. 20, 619–628 (2011).

3. Kwong, W. K. et al. Dynamic microbiome evolution in social bees. Science advances 3, (2017).

4. Koch, H. & Schmid-Hempel, P. Socially transmitted gut microbiota protect bumble bees against an intestinal parasite. Proc. Natl. Acad. Sci. U. S. A. 108, (2011).

5. Martinson, V. G., Magoc, T., Koch, H., Salzberg, S. L. & Moran, N. A. Genomic Features of a Bumble Bee Symbiont Reflect Its Host Environment. Appl. Environ. Microbiol. 80, 3793 (2014).

6. Keller, A. et al. (More than) Hitchhikers through the network: the shared microbiome of bees and flowers. Current opinion in insect science 44, (2021).

7. Zemenick, A. T., Vannette, R. L. & Rosenheim, J. A. Linked networks reveal dual roles of insect dispersal and species sorting for bacterial communities in flowers. Oikos 130, 697–707 (2021).

8. McFrederick, Q. S. et al. Flowers and Wild Megachilid Bees Share Microbes. Microb. Ecol. 73, 188–200 (2017).

9. Müller, H. The Effect of the Change of Colour in the Flowers of ‘Pulmonaria offcinalis’ upon its Fertilisers. Nature 28, 81–81 (1883).

10. Chittka, L., Gumbert, A. & Kunze, J. Foraging dynamics of bumble bees: correlates of movements within and between plant species. Behav. Ecol. 8, 239–249 (1997).

11. Cresswell, J. E. & Robertson, A. W. Discrimination by Pollen-Collecting Bumblebees among Differentially Rewarding Flowers of an Alpine Wildflower, Campanula rotundifolia (Campanulaceae). Oikos 69, 304–308 (1994).

12. Kwong, W. K. et al. Dynamic microbiome evolution in social bees. Science advances 3, (2017).

13. Koch, H., Abrol, D. P., Li, J. & Schmid-Hempel, P. Diversity and evolutionary patterns of bacterial gut associates of corbiculate bees. Mol. Ecol. 22, 2028–2044 (2013).

14. Kwong, W. K., Engel, P., Koch, H. & Moran, N. A. Genomics and host specialization of honey bee and bumble bee gut symbionts. Proc. Natl. Acad. Sci. U. S. A. 111, (2014).

15. Kosior, A. et al. The decline of the bumble bees and cuckoo bees (Hymenoptera: Apidae: Bombini) of Western and Central Europe. Oryx 41, 79–88 (2007).

16. Colla, S. R. & Packer, L. Evidence for decline in eastern North American bumblebees (Hymenoptera: Apidae), with special focus on Bombus affinis Cresson. Biodivers. Conserv. 17, 1379–1391 (2008).

17. Goulson, D., Lye, G. C. & Darvill, B. Decline and conservation of bumble bees. Annu. Rev. Entomol. 53, 191–208 (2008).

18. Williams, P. H. & Osborne, J. L. Bumblebee vulnerability and conservation world-wide. Apidologie 40, 367–387 (2009).

19. Cameron, S. A. et al. Patterns of widespread decline in North American bumble bees. Proc. Natl. Acad. Sci. U. S. A. 108, 662–667 (2011).

20. Goulson, D. Bumblebees: Behaviour, Ecology, and Conservation. (OUP Oxford, 2010).

21. Pascall, D. J., Tinsley, M. C., Clark, B. L., Obbard, D. J. & Wilfert, L. Virus Prevalence and Genetic Diversity Across a Wild Bumblebee Community. Front. Microbiol. 12, 650747 (2021).

22. Brown, M. J. F., Schmid-Hempel, R. & Schmid-Hempel, P. Strong context-dependent virulence in a host–parasite system: reconciling genetic evidence with theory. J. Anim. Ecol. 72, 994–1002 (2003).

23. Shykoff, J. A. & Schmid-Hempel, P. Incidence and effects of four parasites in natural populations of bumble bees in Switzerland. Apidologie 22, 117–125 (1991).

24. Imhoof, B. & Schmid-Hempel, P. Colony success of the bumble bee, Bombus terrestris, in relation to infections by two protozoan parasites, Crithidia bombi and Nosema bombi. Insectes Soc. 46, 233–238 (1999).

25. Averill, A. L., Couto, A. V., Andersen, J. C. & Elkinton, J. S. Parasite Prevalence May Drive the Biotic Impoverishment of New England (USA) Bumble Bee Communities. Insects 12, (2021).

26. Gillespie, S. Factors affecting parasite prevalence among wild bumblebees. Ecol. Entomol. 35, 737–747 (2010).

27. Schmid-Hempel, P. & Schmid-Hempel, R. Transmission of a Pathogen in Bombus terrestris, with a Note on Division of Labour in Social Insects. Behav. Ecol. Sociobiol. 33, 319–327 (1993).

28. Otterstatter, M. C. & Thomson, J. D. Contact networks and transmission of an intestinal pathogen in bumble bee (Bombus impatiens) colonies. Oecologia 154, 411–421 (2007).

29. Durrer, S. & Schmid-Hempel, P. Shared use of flowers leads to horizontal pathogen transmission. Proceedings of the Royal Society of London. Series B: Biological Sciences 258, 299–302 (1997).

30. Brown, M. J. F., Loosli, R. & Schmid-Hempel, P. Condition-dependent expression of virulence in a trypanosome infecting bumblebees. Oikos 91, 421–427 (2000).

31. Martinson, V. G., Moy, J. & Moran, N. A. Establishment of characteristic gut bacteria during development of the honeybee worker. Appl. Environ. Microbiol. 78, 2830–2840 (2012).

32. Ellegaard, K. M. et al. Extensive intra-phylotype diversity in lactobacilli and bifidobacteria from the honeybee gut. BMC Genomics 16, 284 (2015).

33. Kwong, W. K., Steele, M. I. & Moran, N. A. Genome Sequences of Apibacter spp., Gut Symbionts of Asian Honey Bees. Genome Biol. Evol. 10, 1174–1179 (2018).

34. Näpflin, K. & Schmid-Hempel, P. Immune response and gut microbial community structure in bumblebees after microbiota transplants. Proc. Biol. Sci. 283, (2016).

35. Mockler, B. K., Kwong, W. K., Moran, N. A. & Koch, H. Microbiome Structure Influences Infection by the Parasite Crithidia bombi in Bumble Bees. Appl. Environ. Microbiol. 84, (2018).

36. Cariveau, D. P., Elijah Powell, J., Koch, H., Winfree, R. & Moran, N. A. Variation in gut microbial communities and its association with pathogen infection in wild bumble bees (Bombus). ISME J. 8, 2369–2379 (2014).

37. Straw, E. A., Mesnage, R., Brown, M. J. F. & Antoniou, M. N. No impacts of glyphosate or Crithidia bombi, or their combination, on the bumblebee microbiome. Sci. Rep. 13, 8949 (2023).

38. Popp, M., Erler, S. & Lattorff, H. M. G. Seasonal variability of prevalence and occurrence of multiple infections shape the population structure of Crithidia bombi, an intestinal parasite of bumblebees (Bombus spp.). Microbiologyopen 1, 362 (2012).

39. Mallick, H. et al. Multivariable association discovery in population-scale meta-omics studies. PLoS Comput. Biol. 17, e1009442 (2021).

40. Powell, E., Ratnayeke, N. & Moran, N. A. Strain diversity and host specificity in a specialized gut symbiont of honeybees and bumblebees. Mol. Ecol. 25, 4461–4471 (2016).

41. Bosmans, L. et al. Hibernation Leads to Altered Gut Communities in Bumblebee Queens (Bombus terrestris). Insects 9, (2018).

42. Koch, H., Cisarovsky, G. & Schmid-Hempel, P. Ecological effects on gut bacterial communities in wild bumblebee colonies. J. Anim. Ecol. 81, 1202–1210 (2012).

43. Williams, P. H., Thorp, R. W., Richardson, L. L. & Colla, S. R. Bumble Bees of North America: An Identification Guide. (Princeton University Press, 2014).

44. Comeau, A. M., Li, W. K. W., Tremblay, J.-É., Carmack, E. C. & Lovejoy, C. Arctic Ocean Microbial Community Structure before and after the 2007 Record Sea Ice Minimum. PLoS One 6, e27492 (2011).

45. Bolyen, E. et al. Reproducible, interactive, scalable and extensible microbiome data science using QIIME 2. Nat. Biotechnol. 37, 852–857 (2019).

46. Callahan, B. J. et al. DADA2: High-resolution sample inference from Illumina amplicon data. Nat. Methods 13, 581–583 (2016).

47. Rognes, T., Flouri, T., Nichols, B., Quince, C. & Mahé, F. VSEARCH: a versatile open source tool for metagenomics. PeerJ 4, e2584 (2016).

48. Quast, C. et al. The SILVA ribosomal RNA gene database project: improved data processing and web-based tools. Nucleic Acids Res. 41, D590–D596 (2012).

49. Katoh, K. & Standley, D. M. MAFFT multiple sequence alignment software version 7: improvements in performance and usability. Mol. Biol. Evol. 30, 772–780 (2013).

50. Price, M. N., Dehal, P. S. & Arkin, A. P. FastTree 2--approximately maximum-likelihood trees for large alignments. PLoS One 5, e9490 (2010).

51. Cameron, S. A., Hines, H. M. & Williams, P. H. A comprehensive phylogeny of the bumble bees (Bombus). Biol. J. Linn. Soc. Lond. 91, 161–188 (2007).

52. R Core Team. R: A Language and Environment for Statistical Computing. Preprint at https://www.R-project.org/ (2022).

53. Li, D., Dinnage, R., Nell, L. A., Helmus, M. R. & Ives, A. R. phyr: An r package for phylogenetic species-distribution modelling in ecological communities. Methods Ecol. Evol. 11, 1455–1463 (2020).

